# COMPARATIVE ANALYSIS OF TRIPLET COMPOSITION OF COMMON MITOCHONDRIAL AND CHLOROPLAST GENES OF THE SAME SPECIES

**DOI:** 10.1101/2020.12.18.423570

**Authors:** Michael G. Sadovsky, Viktoriya D. Fedotovskaya

**Affiliations:** Institute of computational modelling of SD of RAS; 660036 Russia, Krasnoyarsk, Akademgorodok; V.F.Voino-Yasenetsky Krasnoyarsk state medical university; 660021 Russia, Krasnoyarsk, Partizana Zheleznaka str., 1; Siberian Federal university; 660041 Russia, Krasnoyarsk, Svobodny prosp.79

**Keywords:** frequency, triplet, order, cluster, elastic map, evolution

## Abstract

We studied the relation between the genes encoding the same protein (ATP synthase) in mitochondria and chloroplasts, of the same species. 85 species are studied. The relation is revealed through the unsupervised clustering via elastic map implementation of the points in 64-dimensional space of the triplet frequencies of the genes. The triplet composition was counted with a nucleotide shift of the reading frame along a gene. Three types of clustering have been analyzed: for mitochondria genes solely, for chloroplast genes solely, and for the merged set of the genes from the genomes of both organellae. It was found that the encoded function is the feature in clustering: all the clusters in all three versions of clustering patterns clearly exhibit distinct separation of the genes encoding the same subunit into a separate cluster. This behaviour was found for all three types of cluster patterns.

## I. INTRODUCTION

An interplay between structure of genetic entities, the function encoded in them, and taxonomy of the bearers is a core issue in up-to-date molecular biology and bioinformatics. Evidently, there might be various points of view on this problem, and the answer strongly depends on the specific view. Keeping away from the discussion of the entire variety of the approaches and results obtained in this direction, let us focus on the specific genetic objects and specific structure concept.

To begin with, we fix a structure to be considered everywhere further. We shall investigate the nucleotide sequences of genes common for mitochondria and chloroplasts of the same species; speaking in advance, there are only two groups of such genes: the former are ATP-synthase genes, and the latter are NADH dehydrogenase genes. So, everywhere further a triplet frequency dictionary is stipulated to be a structure of a gene.

Next, we investigate the interplay between two specific parts of a plant genome: these are mitochondria genes vs. chloroplasts ones. Again, one should keep in mind that we study the genes of the same function, in these two genomes. Finally, we will try to trace the impact of the real taxonomy variation on the observed results.

Basically, the approach to investigate the interplay is following. All gene sequences are converted into triplet frequency dictionaries 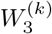. That latter is the list of 64 triplets (lexicography ordered) so that the frequency of the occurrence is indicated for each triplet. Obviously, each dictionary is labeled with (*k*) mark providing the information on the gene type, species, etc. Rigorous statements and exact definitions see below (also see [2, 8, 9, 11] for details).

Conversion of a genetic sequence into a frequency dictionary substitutes a genetic entity with mathematical one: indeed, a frequency dictionary is a point in multi-dimensional metric space (dimension is 63, in the case of triplets). This transform allows to apply the relevant mathematical tools, for a study. The key idea of this paper is to develop a cluster pattern, if any, in 63-dimensional metric space, and check whether a cluster comprises mainly uniformly defined objects: e. g., the genes of the same function, of the genes of the similar genomic origin, etc. The efficiency of such approach has been approved previously (see [1, 2, 10] for details).

Clustering itself is rather complicated problem where the result often depends on the specific details of the methods implemented to do it. Thus, a cluster pattern may not be separated from a method. Here we have used the most powerful and advanced technique of the approximation of multidimensional data with manifolds of small dimension (that is two, to be exact), furthered with clustering provided by local density pattern identification; some essential details of the approach could be found in [3–5].

In brief, the key result of the paper is that the genes corresponding to different genes, both for mitochondria, and for chloroplasts, set up clearly identified clusters so that one cluster comprises one gene. Moreover, the same is true for the case of a combined analysis of the genes of mitochondria and chloroplasts.

## II. MATERIALS AND METHODS

### A. Frequency dictionary

Let now introduce some basic notions and concepts. Let 𝔗 be a sequence from four-letter alphabet ℵ = {A, C, G, T}; biologically, it corresponds to DNA sequence of a gene found in mitochondrion genome, or in chloroplast genome. The length *N* = | 𝔗 | of a sequence is just the number of nucleotides in it.

The triplet frequency dictionary 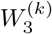 is the list of all 64 triplets *ω* = *ν*_1_*ν*_2_*ν*_3_ counted within a given sequence, with the step *t* = 1 of the reading frame moving right (for certainty) over a sequence. A sequence 𝔗 is connected into a ring, for technical reasons [2].

Everywhere further the dictionaries corresponding to mitochondria genes are shown with triangle labels, while the dictionaries corresponding to chloroplast genes are shown with circles (see Fig. 1).

**FIG 1:**
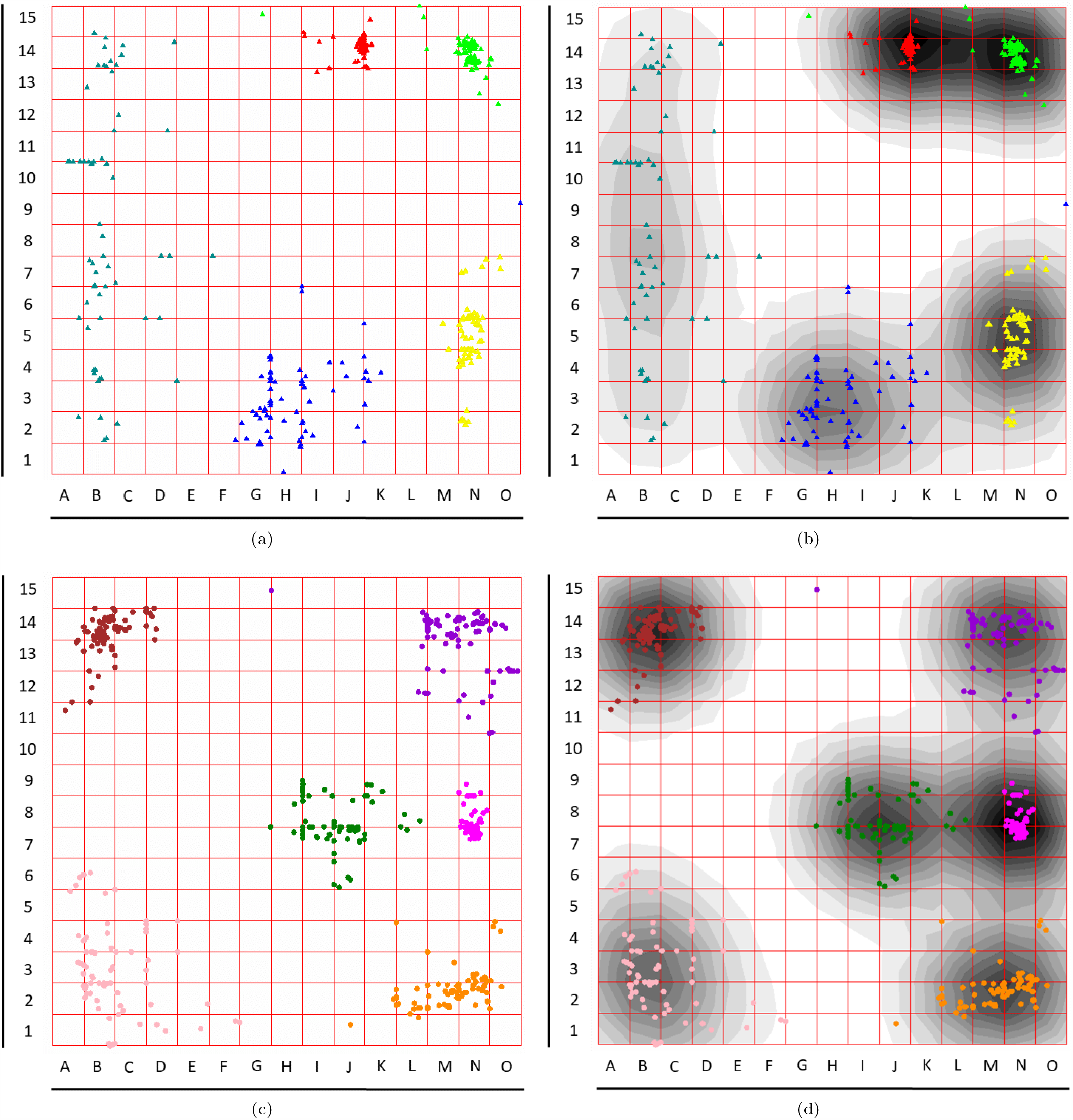
Cluster patterns observed on the set of ATP synthase genes of mitochondria (Figs. 1(a) and 1(b)), and on the set of ATP synthase genes of chloroplasts (Figs. 1(c) and 1(d)). Take a note that mitochondrion genes are labeled as triangles, and chloroplast ones are labeled as circles.

### B. Genetic material

We retrieved genomes of chloroplasts and mitochondria from NCBI bank. The lists of genomes of these two organellae deposited in the bank are rather extended; however, we seek for the organellae genomes belonging to the same species (or even the same specimen, where possible). There are 85 species with both genomes deciphered and deposited in NCBI bank.

These 85 couples of the genomes have been downloaded from the bank, and the reciprocal genes were retrieved from them using CLC WorkBench. A chloroplast genome bears 51 genes, and a mitochondrion genome bears 31 genes, totally. Surely, here we speak on protein coding genes, only. There are two groups of reciprocal genes, in chloroplast genomes and mitochondrion genomes:

− ATP synthase genes, and
− NADH dehydrogenase genes.

It should be stressed that the genes of the same function retrieved from the genomes of different organellae are identical in neither sense. To begin with, mitochondrion ATP synthase genes set consists of five subunits, while that latter from chloroplast genome comprises six subunits. Correspondingly, the sets of genes of NADH dehydrogenase comprise nine and eleven subunits. Here we focus on ATP synthase genes, solely.

### C. Elastic map technique

Elastic map technique is rather novice, so we shall introduce it here in few detail. It implies six steps:

1. find the first and the second principal components, and put on a plane over them (as on axes);
2. project each data point on the plane and connect them to projections with a mathematical spring with infinite expansibility and the elasticity coefficient remains permanent, for any expansion;
3. figure out the minimal square comprising all the projections (so that the rest of the plane must be omitted) and change it with an elastic membrane that is homogeneous: it may bend and expand regardless a direction and/or location;
4. release the system to reach the minimum of the total deformation energy so that elastic membrane transforms into a jammed surface;
5. redefine each point on the jammed surface through the orthogonal projection: i. e. find the nearest point on the jammed surface, for each data point;
6. finally, cut-off all springs to release the jammed surface coming back to a plane.

Clusters were identified through the local density of points. To define it, supply each data point image on an elastic map (in so called inner coordinates, when the jammed surface is already flattened) with a bell-shaped function, e. g. Gaussian one,

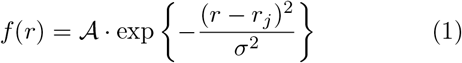

and calculate the sum of them:

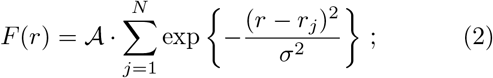

here *r*_*j*_ is the coordinate vector of *j*-th point. Then function *F* (*r*) shows clusters in elastic map. Here *σ* is an adjusting parameter. We used freely distributed software *VidaExpert* to analyze the data. Standard soft 16 ×16 elastic map has been implemented, with by default correlation radius (see eq. (2)) *σ*= 0.25.

## III. RESULTS AND DISCUSSION

Previously, high quality resolution of ATP synthase genes encoding three types of subunits of the protein in fungi mitochondria genomes has been reported [1, 10]. These papers present the clustering probided both by *K*-means and elastic map technique carrie out over the set of the genes encoding three subunits of fungi mitochondrial ATP synthase. Approximately 200 species were involved into analysis; the key result presented in the cited papers consists in the strong prevalence of function over taxonomy. It means that the points corresponding to *W*_3_ dictionaries of the genes set up three clusters, and the clusters comprise the genes of the same subunit, regardless the species.

Here we go further in this direction, and check whether the genes of ATP synthase subunits exhibit similar behaviour, for the genes extracted from other genomes. Moreover, we check the impact of “pseudo-taxonomy”: the difference in the statistical properties of the genes encoding the same function, but extracted from two independent genomes, which are mitochondrion and chloroplast ones. “Pseudo-taxonomy” here means that the genomes are taken from the same species of a plant; in such capacity, they have identical taxonomic position. However, one may expect to face a divergence between them resulted from the different origin of each genome. Here we present the results answering three questions:

- Is it true that the genes of ATP synthase subunits extracted from mitochondrion genome of a plant set up a cluster pattern, and if yes, then is it true that the clusters comprise (mostly) the genes of the same subunit?
- Is it true that the genes of ATP synthase subunits extracted from chloroplast genome of a plant set up a cluster pattern, and if yes, then is it true that the clusters comprise (mostly) the genes of the same subunit?
- Finally, is it true that the genes of ATP synthase sub-units extracted from both mitochondrion and chloroplast genomes of the same plant species set up a cluster pattern, and if yes, then what is the composition of those clusters, in terms of gene origin?

Speaking in advance, the answers on the first and the second questions are positive. Also, the clusters obtained over the database comprising the genes from two genomes simultaneously exhibit very high level of resolution of the genes, both in terms of the encoded subunit, and the genome origin.

Let now explain the coloring scheme used in Fig. 1. All five genes of the mitochondrion ATP synthase subunits are colored in the following manner: *atp*1 is colored in red (RGB scheme is [255, 0, 0]); *atp*4 is colored in lime (RGB scheme is [0, 255, 0]), *atp*6 is colored in blue (RGB scheme is [0, 0, 255]); *atp*8 is colored in yellow (RGB scheme is [255, 255, 0]), and finally, *atp*9 is colored in dark cyan (RGB scheme is [0, 139, 139]).

We used 16 ×16 soft elastic map with by default parameters setting. Everywhere in Figs. 1 and 2 the left sub-figure shows the distribution of the genes over the elastic map in inner coordinates; the right subfigure shows the same distribution with local density level indication. We used 15 levels scheme, elsewhere. To locate a point or a cluster, each map is provided with the chess-like coordinate system.

**FIG 2:**
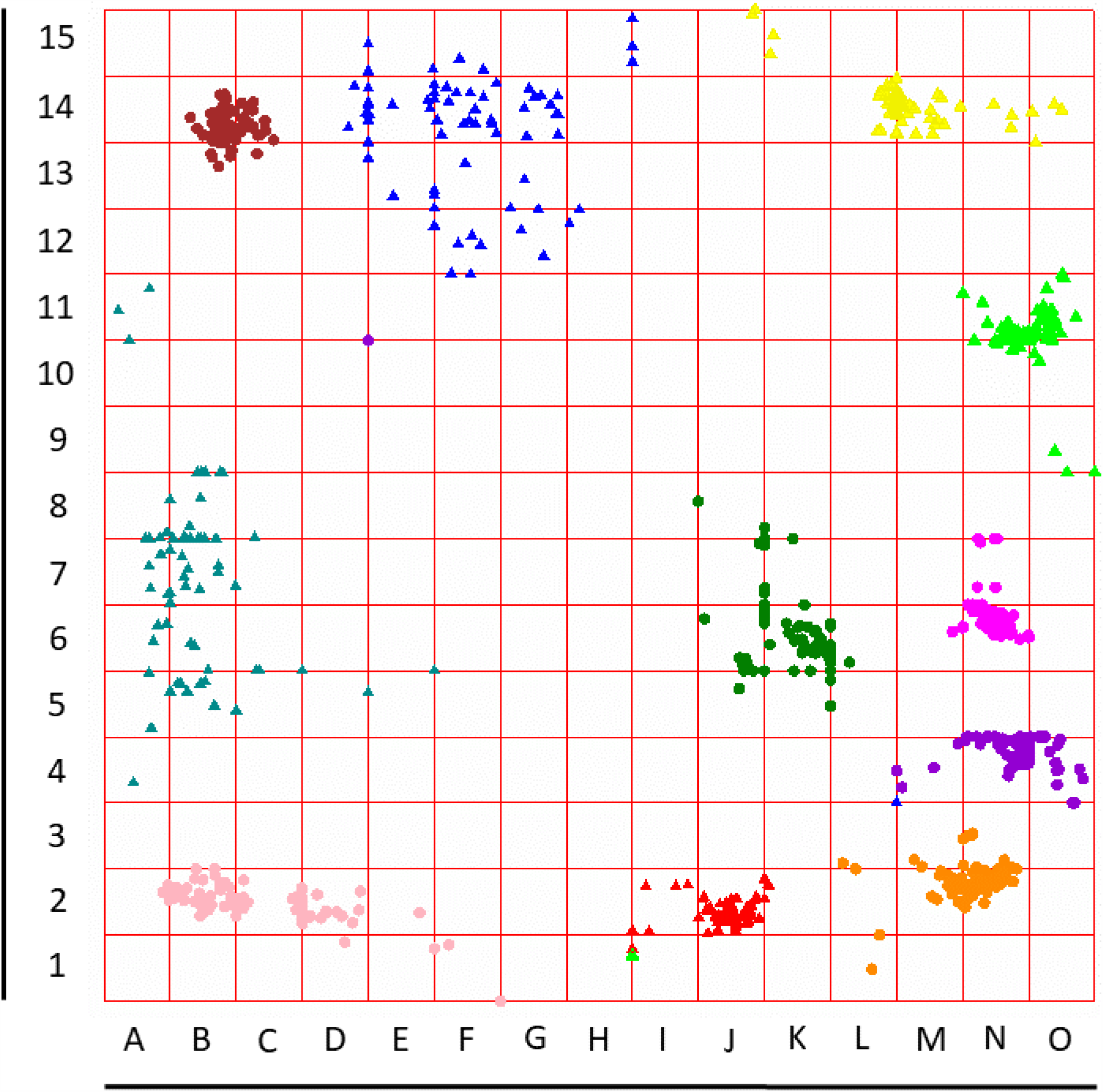
Cluster patterns observed on the set of ATP synthase genes of both mitochondria and chloroplasts. Take a note that mitochondrion genes are labeled by triangles, and chloroplast ones are labeled by circles.

Similarly, six genes of chloroplast ATP synthase sub-units genes are colored in the following manner: *atp*A is colored in magenta (RGB scheme is [255, 0, 255]); *atp*B is colored in green (RGB scheme is [0, 128, 0]), *atp*E is colored in dark orange (RGB scheme is [255, 140, 0]); *atp*F is colored in dark violet (RGB scheme is [148, 0, 211]); *atp*H is colored in light pink (RGB scheme is [255, 182, 193]), and finally, *atp*9 is colored in brown (RGB scheme is [165, 42, 42]).

Let now focus on Fig. 1; Figs. 1(b) and 1(d), to be exact. These figures show the clusters identified through the local density approach. The is an evident difference between these two figures: while chloroplast genes yield six distinctive and unambiguous cluster, thus meeting the number of subunit genes of ATP synthase. The situation is less evident for mitochondria genomes: formally speaking, there only four clusters in Fig. 1(b) identified with local density. We applied the same contrast value *σ* = 0.25, for both maps. So, chloroplast genomes exhibit better distinguishability, when compared to the mitochondrion ones. Obviously, the number of clusters and their occurrence heavily depend on *σ*. Probably, one needs some further studies of the fine structure of the cluster pattern in these two cases. In particular, the observed difference in the clustering is awaiting for the comparative study of the specific genes from mitochondrion genome, and chloroplast genome, to reveal the proximity (or the lack of that latter) of the genes comprised in the tight cluster shown in Fig. 1(b). This study should involve both the genes from the same genome, and the genes from functionally different genomes.

The upper part of Fig. 1 shows the distribution of mitochondria genes, and the lower one shows similar distribution of chloroplast genes. Both subfigures unambiguously exhibit very high level of resolution of the clusters obtained due to elastic map technique: the number of clusters perfectly corresponds to the number of subunits of ATP synthase genes. A distinguishability of the clusters is obvious, for both genetic systems. On the contrary, classification of the genes with *K*-means exhibits rather poor feasibility.

Consider now the joined distribution of the genes of ATP synthase subunits gathered from both genetic systems (see Fig. 3); here the same labeling and coloring schemes are applied. First of all, one can see just seven clusters, if identified through the local density (see Eqs. (1, 2); these are [B – D, 2], [B, 5 – 8], [B – E, 13 – 14], [M, 14], [N, 10 – 11], [J, 2], [K, 6], and, finally, [M – N, 2 – 6] (the largest one). Note, we discuss the cluster pattern revealed through local density (1, 2) implementation; other methods (e. g., graph theory based ones [6, 7, 12–15]) may yield different cluster patterns.

**FIG 3:**
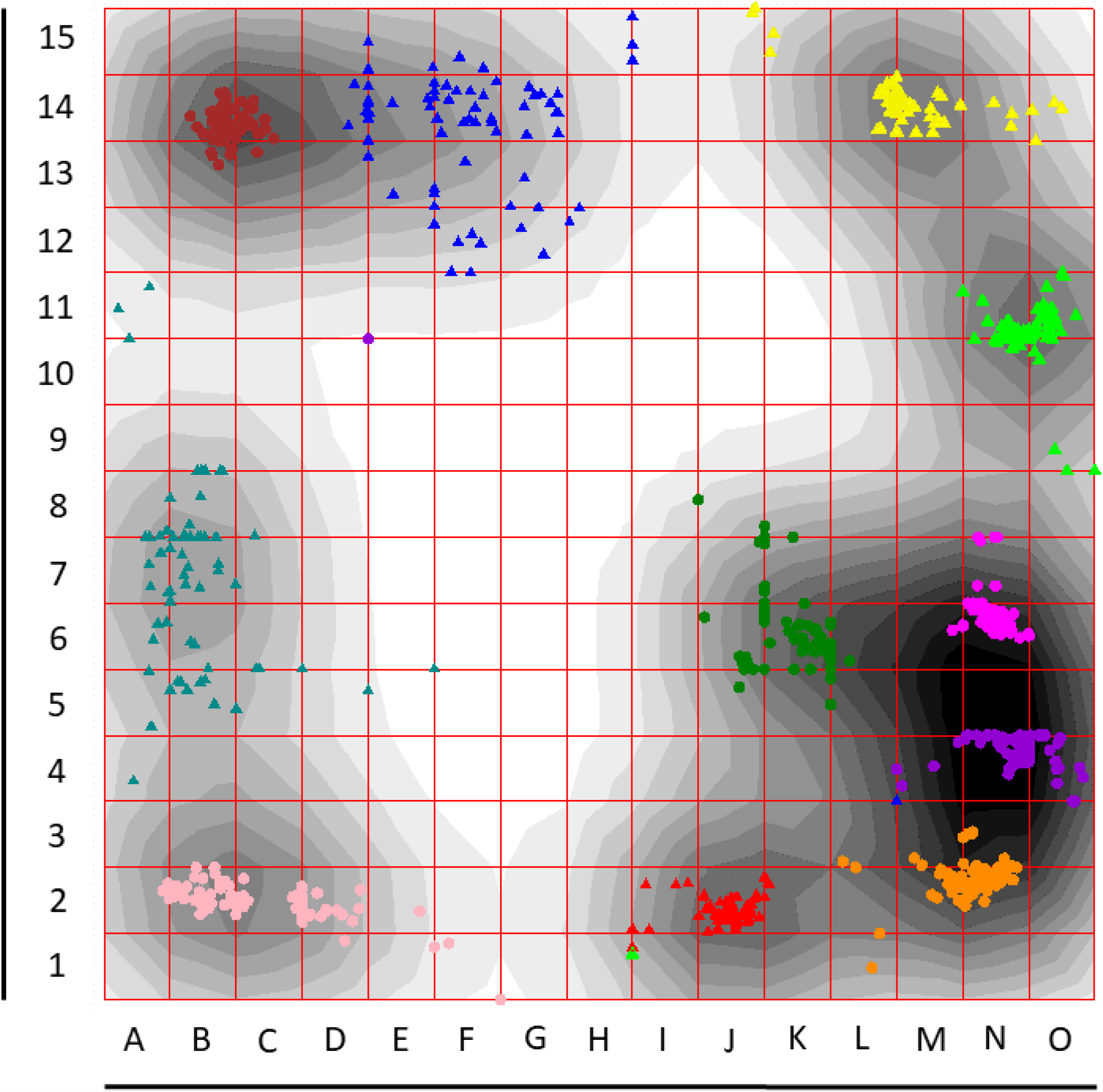
Cluster patterns observed on the set of ATP synthase genes of both mitochondria and chloroplasts, with local density indicated on map. Take a note that mitochondrion genes are labeled by triangles, and chloroplast ones are labeled by circles.

Local density algorithm may mimicries a cluster pattern: indeed, even a fast glance on Fig. 2 unambiguously reveals eleven clusters comprising the genes of reciprocal subunits, separately for mitochondria, and for chloroplasts. First of all, one sees the highest specificity of the clusters: there are very few point with improper cluster location. Indeed, these are just three genes outlying their “home” cluster: *atp*4 by *Welwitshia mirabilis* ([H I, 1]), and the couple *atp*6 by *Ammopiptanthus mongolicus* – *atp*F by *Ammopiptanthus nanus* ([L – M, 3 – 4] and ([D – E, 10 – 11])) where the genes change their regular cluster occupation.

Chloroplast genomes differ mitochondria ones from the point of view of a dispersion of a cluster: the chloroplast genomes form very tight clusters, with a single exclusion, that is *atp*H cluster ([B – D, 2]). This cluster is split into two distinct subclusters, and the reason for such behaviour still awaits for investigation. On the contrary, the clusters of *atp*6 and *atp*9 subunits genes exhibit rather disperse pattern; at least, the obviously differ from any other one, from that point of view.

## IV. CONCLUSION

Here we investigated the interplay between the structure of nucleotide sequences of specific genes, the function encoded in them, and the origin of a gene. A structure may be defined in various ways; however, we have used probably the simplest one, that is the triplet frequency dictionary. An intention to take into consideration the triplet composition is quite clear: the triplets play exceptional role in inherited biological information processing.

To compare the impact of an origin of genes taken into consideration was the key idea of our research; indeed, we used the genes of the same function (that is ATP synthase set of genes), taken from the same organism that is a plant. The origin of the genes was the only influential factor in that case: we have taken the genes of ATP synthase from mitochondrion of a plant, and from chloroplast of the same plant. The key question was what is stronger:

- triplet composition proximity;
- similarity of the encoded function, or
- origin of a gene (mitochondrion of chloroplast).

Unsupervised cluster techniques have been implemented to answer the question.

Function encoded in a gene is the strongest among the issues enlisted above. Indeed, the clustering is provided by the triplet composition proximity of gene nucleotide sequences. That is a common place that proximal structure (triplet composition, in our case) follows in a proximity of the encoded functions. An inverse statement is less evident: there is no guarantee that proximal (or even the same) function may not be encoded with significantly different structures. Here we see the duality of these two issues: on one hand, clustering carried out over the triplet composition yields the clusters comprising functionally the same genes; on the other hand, there are very few large deviations of the points from “home” cluster. In fact, there are only three genes exhibiting diverse occupation.

Organella genome origin may be considered as a kind of taxonomy simulation: one may expect that the distribution of the genes encoding the same function are sensitive to the origin, or insensitive. That latter case means that a cluster must comprise the genes from both genomes. Here we see a direct and unambiguous proof of the significant divergence of the genes extracted from different genomes, in the cluster pattern.

